# Time-series monitoring of transgenic maize seedlings phenotyping exhibiting glyphosate tolerance

**DOI:** 10.1101/2022.03.21.485126

**Authors:** Mingzhu Tao, Xuping Feng, Yong He, Jinnuo Zhang, Xiulin Bai, Guofeng Yang, Yuzhen Wei

## Abstract

Developing herbicide resistant cultivars is one of the effective methods to solve the safety problem caused by the use of herbicide. In this study, hyperspectral image was used to develop more robust leaf chlorophyll content (LCC) prediction model based on different datasets to finally analyze the response of LCC to glyphosate-stress. Chlorophyll a fluorescence (ChlF) was used to dynamically monitor the photosynthetic physiological response of transgenic glyphosate-resistant and wild glyphosate–sensitive maize seedlings, and applying chemometrics methods to extract time-series features to screen resistant cultivars. Both the proposed two transfer strategies achieved the best prediction of LCC with a coefficient of determination value of 0.84, and relative root mean square error of 4.03 for the prediction set. Based on the predicted LCC and ChlF data, we found the antioxidant system of glyphosate-sensitive plants is too fragile to protect themselves from the damage, while glyphosate-resistant plants could overcome it by activating more powerful antioxidant system. φ_E0_, V_J_, ψ_E0_, and M_0_ could be used to indicate damage caused by glyphosate to differentiate resistant cultivars. This study provided a new methodology to monitor LCC to finally analyze glyphosate tolerance in a time-series manner, and verified the feasibility of ChlF in crop breeding.

**Highlight:** This study proposed a new methodology to monitor leaf chlorophyll content to finally analyze glyphosate tolerance in vivo, and verified the feasibility of chlorophyll a fluorescence in crop breeding.

## Introduction

Glyphosate is one of the most popular herbicides in grass control with non-selective systemic and broad-spectrum characteristics (Duke and Powles, 2008; de Freitas-Silva *et al.*, 2020). Under high selective pressure of glyphosate, plants are prone to genetic mutations that show tolerance to glyphosate (Duke and Powles, 2008; Beckie, 2011). Simultaneously, it also provides a new way to acquire glyphosate-resistant varieties. With the rapid development of genetic engineering and genomics, the development and popularization of superior varieties can be achieved. Glyphosate has become more ubiquitously used in agriculture since the introduction of glyphosate-resistant cultivars.

In the process of cultivar identifying, the professional technologies, such as PCR, often require expensive experimental equipment and professional operating knowledge, which limits the convenience and real-time detection. Traditional detection method of herbicide resistance is to observe seedlings’ growing status with the naked eye after spraying the herbicide to them. Generally speaking, it takes 12 days from the application of stress to the determination of herbicide injuries level and final resistance identification (Singh *et al.*, 2021), which is time-consuming, subjective and inefficient. Furthermore, biomass, chlorophyll content, and the activity of antioxidant enzyme were often used to evaluate the level of tolerance to herbicide stress of crops. As we know, the photosynthetic pigment is located in the membrane of plant thylakoids that captures and transfers light energy or causes initial photochemical reactions. Physiological requirements and material accumulation capacity of plants can be known according to changes in pigment content (Bloem *et al.*, 2020). Hence, the pigment content in leaves is one of the most frequently-used in the investigation of plant nutrition, health status, and physiological response to environmental conditions including various stresses (Vahtmäe *et al.*, 2018).

Up to this point, spectrophotometer is still a popular method for the determination of photosynthetic pigment content, which is not only destructive, but also difficult to meet the requirements of modern agriculture due to its lag. The emerging hyperspectral technology has yielded a positive foundation in the evaluation of pigment, usually based on machine learning and chemometrics, but the method is still in development stage, and rarely combined its results with specific experimental phenomenon to illustrate the stress response, such as the mechanism of action of glyphosate stress in plants.

As another technique for indirectly sensing plants’ pigment statues, rapid Chl a fluorescence (ChlF) can obtain enough information about the structure and function of photosystem II (PSII) in just a few seconds, and ultimately apply to screen genotype. In recent years, this technology has been ubiquitously adopted to assess the direct effects of biotic and abiotic stresses, such as high temperature (Chen *et al.*, 2016), drought and heavy metals (Stirbet *et al.*, 2018), on photosynthetic physiology of plants. ChlF can detect the stress response before plants exhibit visible signs or other detection methods, such as measurements of gas exchange or changes in pigment (Kuckenberg *et al.*, 2009; Kalaji *et al.*, 2018b). Glyphosate is a kind of abiotic stress that effects the photosynthetic physiological mechanism of plants. There are two target sites in plant chloroplasts. The one is the electron transport chain, which is related to the utilization of absorbed light energy. The other one is the synthesis of Chl and carotenoids, which is associated with light-harvesting complex (LHC) and the antenna of the photosynthetic reaction center (Kalaji *et al.*, 2018b). The above changes can be captured by OJIP curve and JIP-test parameters. ChlF contains the relationship between the light reaction stage and PSII fluorescence (Zhang *et al.*, 2021a), and JIP-test parameters include absorption and trap of energy and electron transport information.

Apart from these, to our knowledge, it has been less studied that applying hyperspectral technology and prompt ChlF transient to dynamically monitor various glyphosate response of maize seedlings in time-series. This study will provide a new rapid phenotypic analysis method for gene function studies, and has the potential to advance the crop breeding. In turn, the gap between phenotype monitor and transgenic technology promotes us to explore the response and characteristic fingerprint of different resistant varieties under glyphosate stress. Therefore, this study aims to (1) develop more robust leaf chlorophyll content (LCC) prediction model; (2) explore the feasibility of rapid ChlF transient to dynamically monitor the photosynthetic physiological response of different resistant maize varieties under glyphosate stress, (3) analyze and find out the effects of glyphosate-stress on plant photosynthetic characteristics, and (4) select characteristic fingerprints to evaluate the tolerance of glyphosate. The results of this study will provide a new tool for the identification of excellent traits which may possibly improve the efficiency of crop breeding in the long run.

## Materials and methods

### Plant materials and experimental setup

Accession of maize including glyphosate-sensitive wild type (recorded as GS) and glyphosate-resistant type (recorded as GR) were provided by the Institute of Insect Sciences, Zhejiang University, Hangzhou, China. Glyphosate-tolerance of GR was acquired by expression of a mutant 5-enolpyruvylshikimate-3-phosphate synthase (EPSPS) enzyme, specific protocol was described in previous research (Feng *et al.*, 2017).

The experiment was conducted in the greenhouse at Zhejiang University. Some parameters followed the published research (Kalaji *et al.*, 2014) with minor modifications. The temperature and photoperiod of day/night were 28/26□ and 11/13 h, respectively, the average relative humidity was adjusted to 55%, and the maximum photosynthetically active radiation was about 1400 μmol (photons) m^-2^ s^-1^. Specifically, maize seeds were sown in seeding clods one per pot, and on the 9^th^ day post sowing (two-leaf stage, the second leaf was fully expanded but the third leaf had just emerged), plants of similar growing state were picked and transplanted into the plastic bucket with drainage holes, each pot was only occupied by one plant. When plants grew to the 3-leaf stage (the third leaf was fully expanded but the fourth leaf had just emerged) on the 15^th^ post sowing, part of plants were subjected to glyphosate (Roundup Original®, Monsanto Company) as treatment at the recommended dose by using portable sprayer, and the others were sprayed with water as control. Therefore, there were four groups (two varieties × two treatments) in our research, namely RT, RW, ST, and SW (“R” stands for “glyphosate-resistant”, “S” stands for “glyphosate-sensitive”, “T” stands for “with glyphosate”, “W” stands for “with water”).

ChlF and hyperspectral measurement were performed at 2, 4, 6, 8 days after treatment (DAT) to monitor and assess photosynthetic apparatus in time scale. No less than four plants of each group at each sampling point were randomly selected, and all the measurements were conducted on the 1^st^, 2^nd^ and 3^rd^ leaves (recorded as L1, L2, L3 from the bottom to top, respectively) of maize plants. The details of each independent experiment were shown in Fig. 1.

**Fig. 1.**
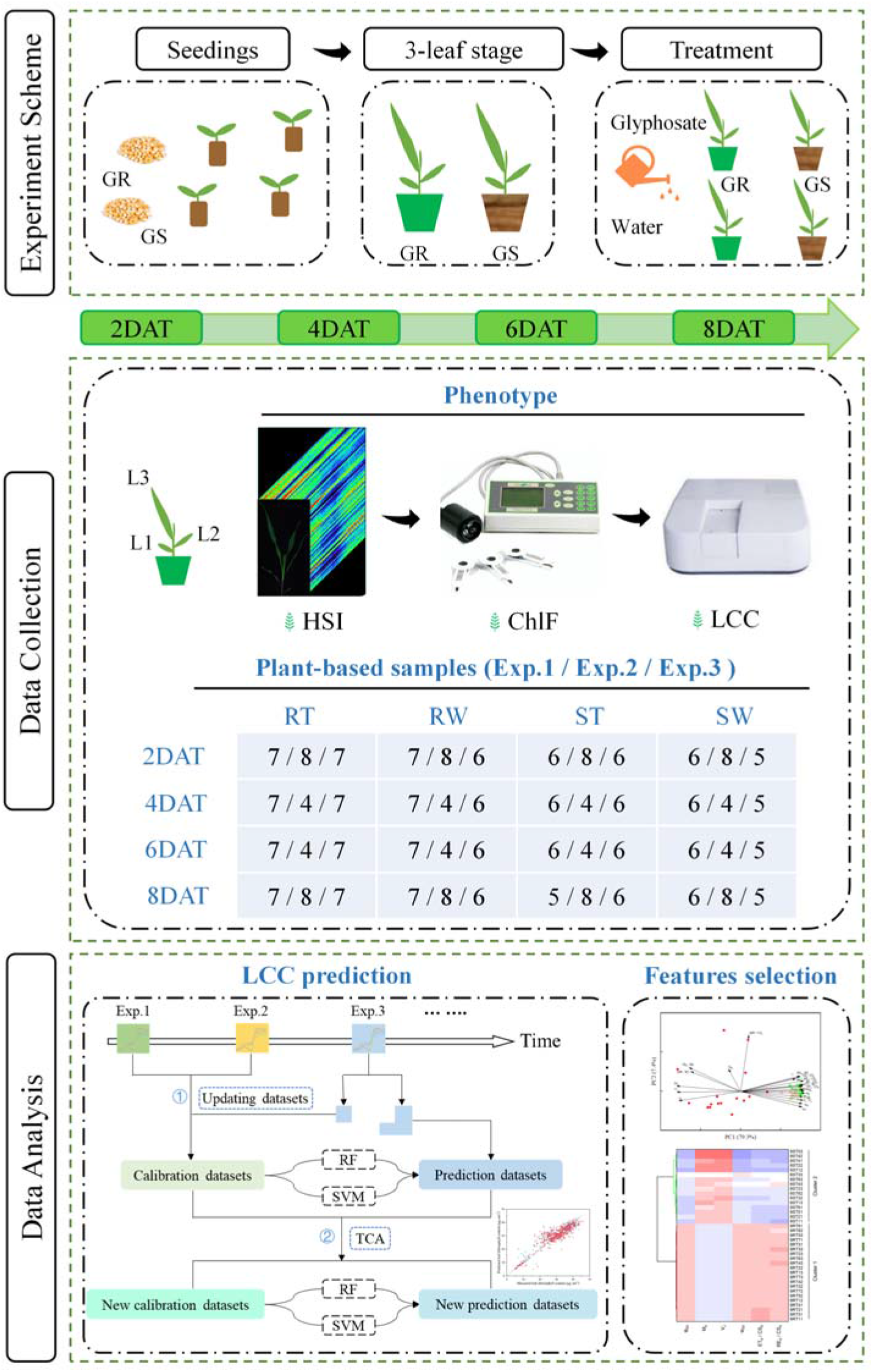
Flowchart of maize phenotype monitoring. Three individual experiments (Exp.1-3) were conducted in the same greenhouse in August, November and December 2021, respectively. GR: glyphosate-resistant cultivar; GS: glyphosate-sensitive cultivar; DAT: days after treatment; HSI: hyperspectral image; ChlF: chlorophyll a fluorescence; LCC: leaf chlorophyll content; RT, RW, ST, and SW: “R” stands for “glyphosate-resistant”, “S” stands for “glyphosate-sensitive”, “T” stands for “with glyphosate”, and “W” stands for “with water”; RF: random forest; SVM: support vector machine; TCA: transfer component analysis.

### Acquisition of hyperspectral imaging in vivo

Hyperspectral image acquisition of fresh maize seedlings was conducted by a line-scan hyperspectral imaging system in the Vis-NIR range (380-1030 nm), which was described in our previous study at length (Feng *et al.*, 2018). Try to keep maize seedlings’ growth morphology in natural and lay them flat on a black plate without overlapping between leaves. In the pre-experiment, in order to guarantee hyperspectral images sharpness and without distortion, we adjusted the camera exposure time, the intensity of line light source, the distance between the camera lens and the moving plate, and the speed of the conveyer belt to 70 ms, 240, 390 mm, and 5 mm/s, respectively.

The original hyperspectral images of the maize seedlings were corrected by the black and write reference images, which were obtained under the same system parameters as samples. Prior to estimating LCC, image segmentation must be applied to identify the entire area of each expanding leaf as the region of interest (ROI). Hence, each plant had three ROIs corresponding to three leaf positions (L1, L2, and L3). For each ROI, after the head-to-tail spectra bands with high noise were discarded, the spectral reflectance of all pixels at 450-902 nm were averaged to represent the corresponding leaf.

### Measurement of ChlF in vivo

Measurement of ChlF transient curves was completed with a continuous excitation chlorophyll fluorimeter (Handy-PEA, Plant Efficiency Analyzer, Hansatech Instruments Ltd., Norfolk, United Kingdom) at room temperature (ca. 26□) from 11:30 to 15:00. After hyperspectral image acquisition, three clips were immediately placed on the basal, central, and top positions of the adaxial side of each leaf, respectively, for a 25 minutes’ dark adaption to allow all PSII reaction centers (RCs) open (Q_A_^-^ gets reoxidized), and ferredoxin-NADP^+^-reductase to inactive. Then a polyphasic ChlF transient curve could be obtained with the illumination of a saturating light excited by an array of three ultra-bright red LED (3000 μmol-m^-2^-s^-1^; peak wavelength at 650 nm). The typical OJIP transient was recorded during the 1 s period that included four steps O (at 20 μs), J (at 2 ms), I (at 30 ms) and P (peak). Once ChlF rise kinetics was obtained, all leaves were stored in zip-lock bags at −80□ until LCC measurement.

### A brief information of OJIP fluorescence transient and JIP-test

OJIP fluorescence transient curve reflects the sequential events with four steps generally (Neubauer and Schreiber, 1987). O-step refers to the initial fluorescence level (*F*_0_). P-step is the peak level with all RCs are closed. J-step and I-step are transitory steady-states of Q_A_^-^ concentration, as the slope of curve at these steps are zero, which means that the reduction rate of Q_A_ is equal to the oxidation rate of Q_A_^-^. Between the two adjacent steps, denoted as phase, reflects different photochemical stages. OJ-phase is mostly driven by primary photochemistry, almost only single-turnover (Q_A_ to Q_A_^-^) events are going on. Because Q_B_ could not accept electron from Q_A_^-^ in time to reoxidized it (the electron transport time is ~250 ps from pheo^-^ to Q_A_, but 0.1~0.6 ms from Q_A_^-^ to Q_B_), resulting in a large accumulation of Q_A_, and causing the fluorescence intensity to increase. JI-phase mainly refers to the reflection of the intersystem electron carriers Q_B_, plastoquinone (PQ), and cytochrome (Cyt b6/f). IP-phase refers to PSI acceptor side (Tsimilli-Michael, 2020).

On the basic of OJIP trajectory and the theory of “energy flow” across thylakoid membranes, quantitative method so-called JIP-test, is used to explore detailed information of PSII status and function including RCs, light-harvesting antenna, and both the donor and acceptor sides. All the JIP-test parameters can be divided into four groups in general: (1) basic measured and calculated values; (2) quantum yield and probabilities/efficiencies; (3) energy flux; (4) performance indices. Among them, the parameters used in this study are listed in detail in Table 1 (Spyroglou *et al.*, 2021) with minor modifications.

**Table 1.**
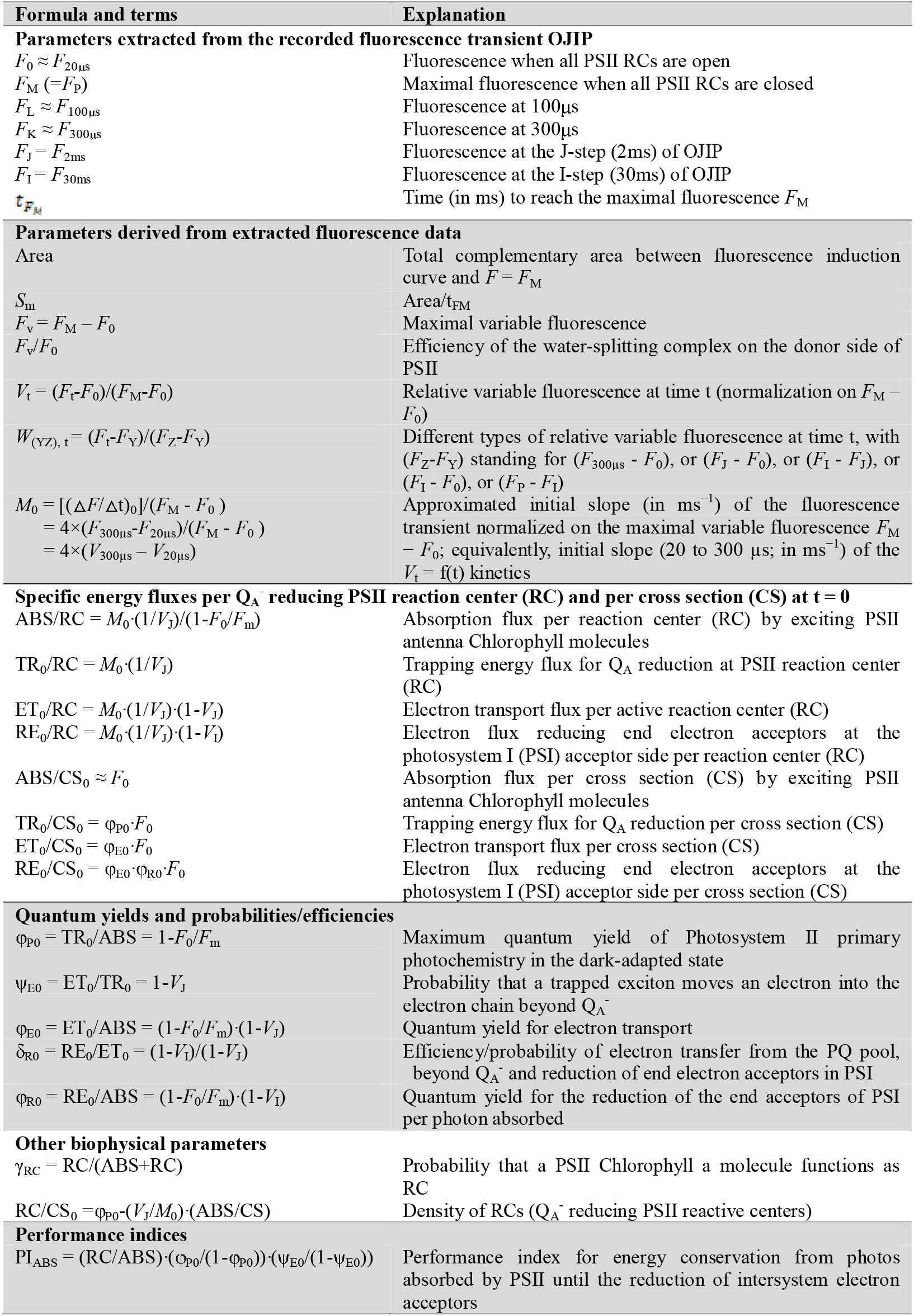
Glossary, formula and explanation of terms used in the JIP-test for analysis OJIP fluorescence transient curve emitted by dark-adapted photosynthetic samples in this study. Subscript “0” (or “o”) indicates that the parameter refers to the onset of illumination. Symbol “*F*” stands for “fluorescence”, means fluorescence intensity.

### Measurement of leaf chlorophyll content

The measurement of LCC was followed the traditional protocol with minor modification (Zhang *et al.*, 2021b). Similar to ChlF acquisition, three discs of 5 mm diameter were taken from the basal, central, and top positions of each leaf, respectively. LCC in discs were extracted entirely after soaking in 4 mL ethanol: water 95% (*v*/*v*) for 48 h in dark environment. Then pigment content was measured by spectrophotometry (Epoch, BioTek Instruments Winooski, United States). Finally, according to the standard curves and the absorbance of light at 470, 649 and 665 nm, Chlorophyll content was calculated. The basal, central, and top positions of each leaf were averaged as the corresponding phenotypic traits of each leaf.

### Data analysis and model development

In this study, based on spectral reflectance datasets from three independent experiments (Exp. 1-3) conducted at different times, we explored the reliability of using data sets obtained at different times as training set and test set, and evaluated the performance of two optimize strategies. In order to overcome experiment condition diversity and improve the robustness of LCC estimation model, the following two strategies (Fig. 1) were used. One was to exploit partial dataset (≤ 50%) of Exp.3 to rebuild training dataset, which were selected by the Kennard-Stone (KS) algorithm. The other strategy is to narrow the distribution difference between source domain and target domain through transfer component analysis (TCA). The above strategies could be regarded as the preprocessing step. To construct prediction, the commonly used random forest (RF) and support vector machine (SVM) regression were employed with the raw datasets (Exp.1 and Exp.2) or updated datasets (adding partial samples from Exp.3 or using TCA algorithm). For RF regression model, the range of the minimum number of observations per tree leaf was set from 5 to 25 with a step of 5, the range of the number of decision trees was set from 100 to 300 with a step of 50, and the optimal value was determined by 10-fold cross validation. For SVM regression model, the epsilon-SVR algorithm was selected to develop the regression model. The epsilon in loss function (p) was set 0.1, and the particle swarm optimization method was applied to optimize model parameters (‘g’: gamma, ranges from 0.1 to 100, ‘c’: cost, ranges from 0.01 to 1000). The performances of prediction models were quantified by the coefficient of determination (R^2^) and the root of mean square error (RMSE). The above algorithms were conducted on Matlab 2016a (The MathWorks, Natick, MA, United States).

With glyphosate applied, plants exhibited complex physiological responses, which subtly varied depending on plant tolerance degree and stress time. To obtain a general view of dataset and understand the resistant mechanism of GR and GS on the time scale, principal component analysis (PCA) was performed to visualize time-series phenotypic data and further dissection intuitively. Analysis of variance (ANOVA) was then applied to quantify the differences of JIP-test parameters between four groups on 2, 4, 6, 8 DAT. Significant differences were evaluated at a significance level (*P*) of 0.05 and Holm-Bonferroni test was used as the *post-hoc* test. As an unsupervised learning method, PCA could replace the original variables with fewer new variables through linear combination, which retained as much information as possible from the original variable. Hence, according to JIP-test parameters’ absolute value, PCA-loadings was applied to select optimal features on each sampling point. Then, the Ward.D linkage clustering method was used to validate the performance of the selected features. The above methods were implemented on Origin 2021b (OriginLab Corporation, Northampton, MA, United States).

## Results

### Establishment of LCC prediction model based on spectral reflectance

Different machine learning algorithms and optimization strategies were employed to predict LCC based on the multi-temporal spectral reflectance, and the results with R^2^, RMSE were shown in Table S1. When spectral datasets of Exp.1 and Exp.2 were used as a calibration set to estimate LCC of Exp.3, the better prediction result was based on the SVM regression model, and obtained R^2^_p_ was 0.65, and RMSE_p_ was 4.94. With adding new samples from Exp.3, there was an increasing tendency of R^2^_p_. In the SVM-based algorithm, the range of R^2^_p_ and RMSE_p_ was 0.70 ~ 0.81, 4.39 ~ 4.80, respectively. In the RF-based algorithm, the range of R^2^_p_ and RMSE_p_ was 0.57 ~ 0.62, 6.11 ~ 6.42, respectively. The best estimation result was found in the SVM regression model when adding 144 new samples from Exp.3 to the first two experiments (Exp.1 and Exp.2) dataset (R^2^_p_ = 0.81, and RMSE_p_ = 4.42). After applying TCA, the estimation results were further improved, which was embodied in the increasing tendency of R^2^_p_ and the decreasing of RMSE. The above indicated that both adding new samples and applying TCA algorithm could narrow the difference between source domain dataset and target domain dataset, and improve prediction performance (Feudale *et al.*, 2002; Weng *et al.*, 2018). By comparing the results of different strategies, TCA_SVM regression model with 144 new samples exhibited the best results (R^2^_p_ = 0.84, and RMSE_p_ = 4.03), which was presented in Fig. 2.

**Fig. 2.**
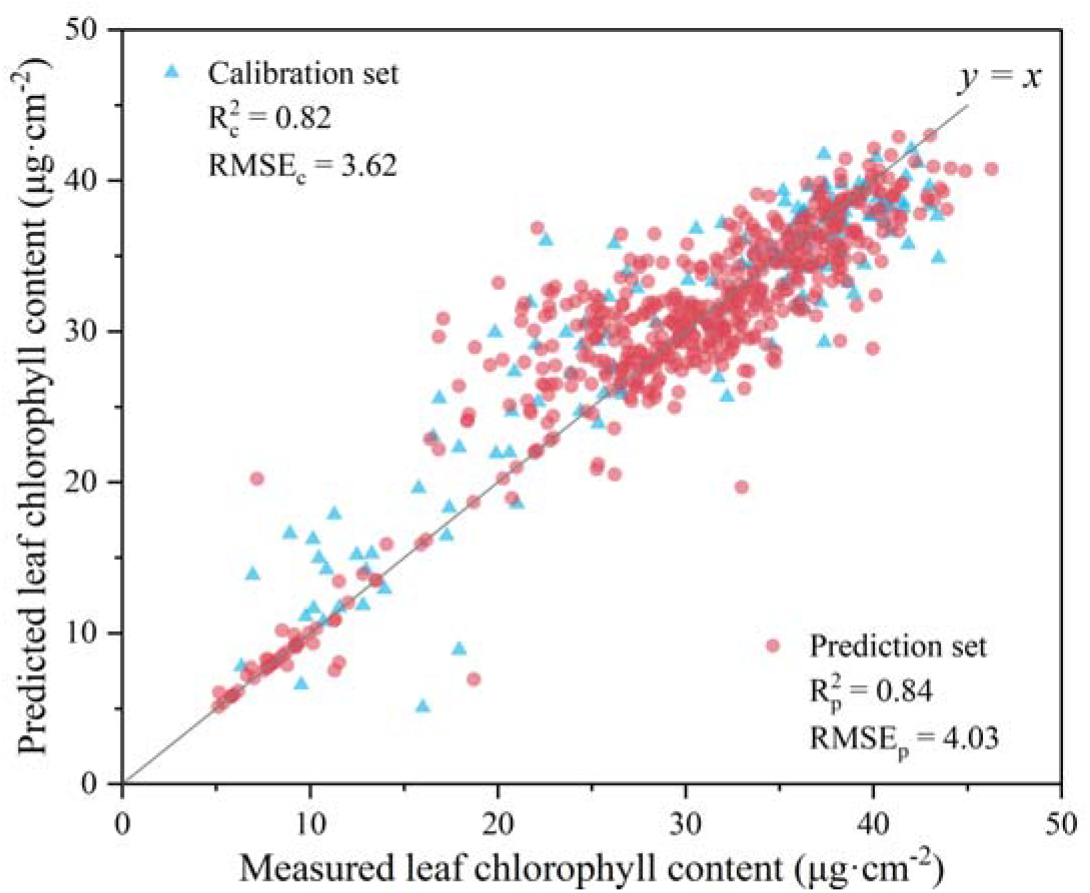
The prediction results of leaf chlorophyll content. The results were accomplished by support vector machine (SVM) regression model developed from multi-temporal spectral reflectance dataset (Exp.1 and Exp.2) with adding an external dataset from Exp.3 using transfer component analysis (TCA). The subscripts c and p represented calibration set and prediction dataset.

### Effect of glyphosate stress on pigment content within leaves of maize seedlings

Based on LCC predicted by the model with the best performance, its ANOVA result was shown in Fig. 3. Although RT plants’ LCC decreased on 2 DAT, there did not show significant difference between RT and RW plants throughout the trial. However, ST plants revealed notable lower LCC when compared to SW on 4 DAT, and this tendency was more obvious on 6 and 8 DAT. This result was similar with previous literatures (Fan *et al.*, 2017; Feng *et al.*, 2018), which indicated that our model can be applied to the detection of physiological indicators without tedious laboratory tests.

**Fig. 3.**
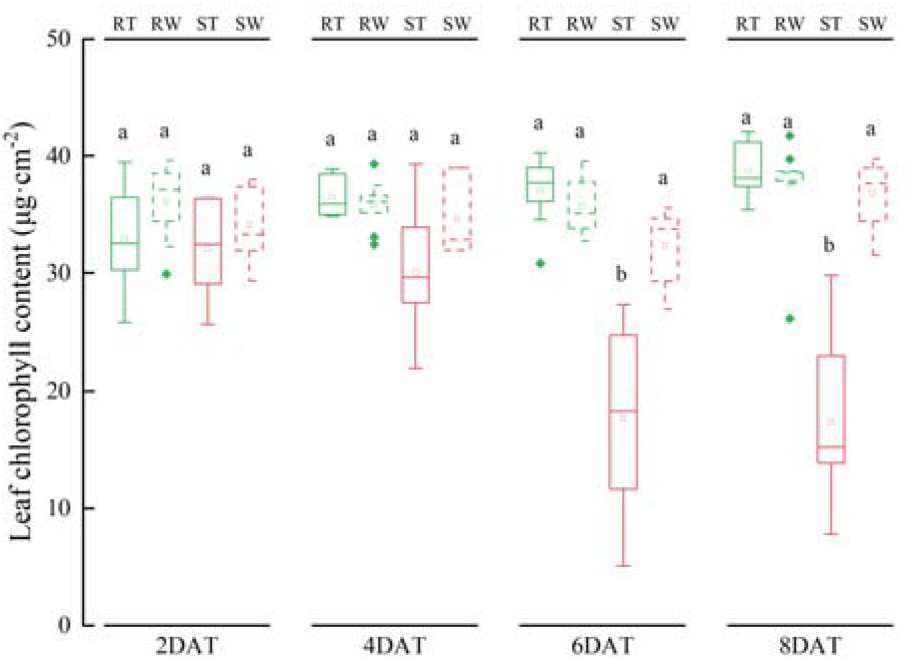
Impact of glyphosate stress on leaf chlorophyll content on 2, 4, 6, and 8 days after treatment (DAT). Within the same sampling point, any two bars with a common letter are not statistically significant according to ANOVA (Holm-Bonferroni post-hoc test, *P* > 0.05). RT, RW, ST, and SW plants are represented by green solid, green dashed, red solid, and red dashed line.

### OJIP transient captures the difference responses to glyphosate stress

The OJIP transient curves of all four groups presented a typical polyphasic O-J-I-P shape on a logarithmic time scale in the early stage of treatment (Fig. 4). The OJIP curves of RW and SW plants were basically the same at four sampling points. This indicated that there was no significant difference in photosynthetic between the two cultivars, and it was difficult to select glyphosate-resistant cultivars by ChlF without treatment. Sprayed with glyphosate did not change the F0 but change fluorescence intensity at J, I, and P steps before the plants died. Regardless of cultivar, P-step was the most sensitive to glyphosate among the four steps in OJIP curves. The fluorescence intensity at P-step (*F*_M_) decreased significantly in both cultivars at 2 DAT. Interestingly, the fluorescence intensity difference between RT and RW decreased from 2 DAT progressively, and at 6 DAT, the difference almost disappeared, which suggested GR plants returned to the normal photosynthetic state at 6 DAT. However, the OJIP curve trend of ST was significantly different from SW at 6 DAT without recovery sign. Therefore, under glyphosate stress, the difference of OJIP curves between GR and GS became obvious with time both in fluorescence intensity and curve morphology, which indicated the feasibility of ChlF in identifying glyphosate-resistant cultivars.

**Fig. 4.**
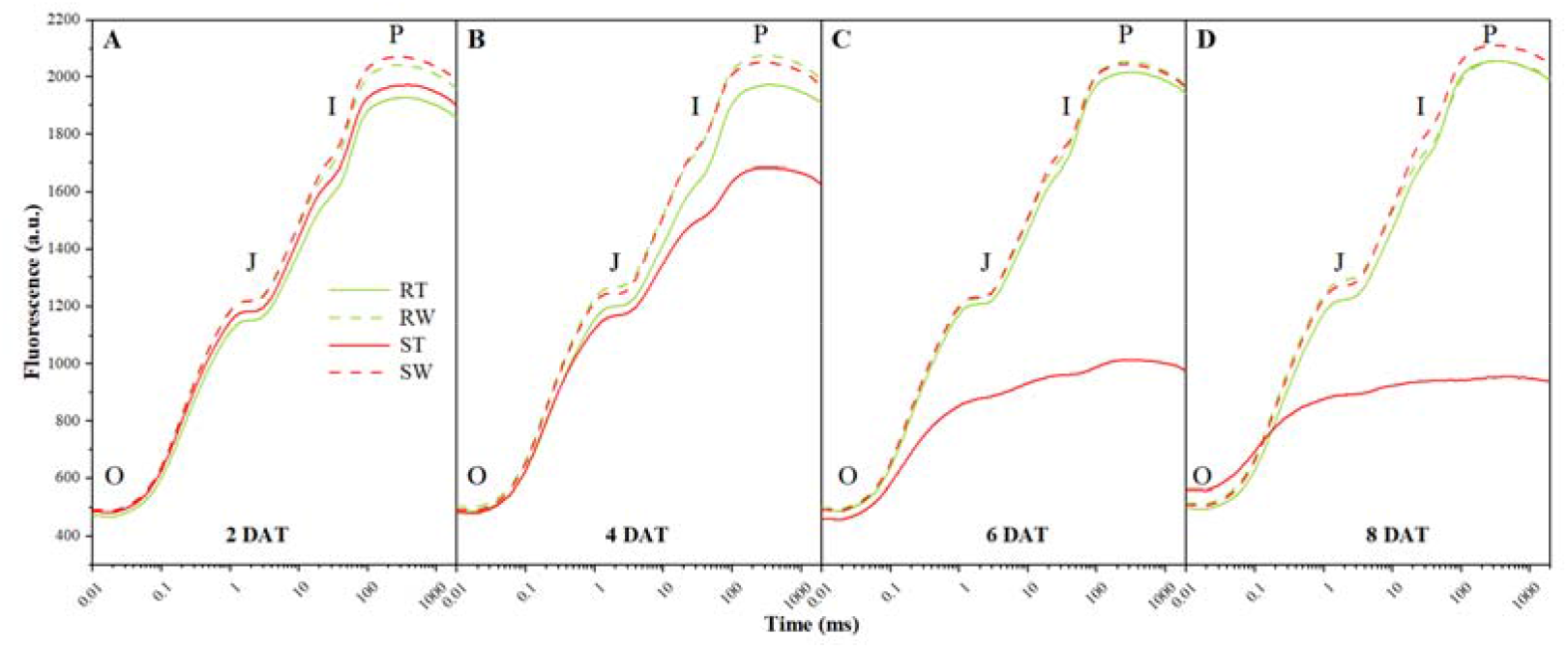
The OJIP transient curve of two maize cultivars (glyphosate-sensitive and glyphosate-resistant) with two treatments (glyphosate and water) at (A) 2, (B) 4, (C) 6, (D) 8 days after treatment (DAT). As for the marks of the four groups, “R” stands for “glyphosate-resistant”, “S” stands for “glyphosate-sensitive”, “T” stands for “with glyphosate”, “W” stands for “with water”.

To further visualize the information hidden in the raw OJIP transient curves, a semiquantitative analysis of normalization (Kalaji *et al.*, 2016) was applied for the better depiction of respond difference between GR and GS. Normalization and corresponding subtractions were done in the OK, OJ, OI, and IP-phase, respectively. The fluorescence rise kinetics was normalized between O-step (20 μs) and K-step (300 μs), as *W*_OK_ = (*F*_t_ – *F*_0_)/(*F*_K_ – *F*_0_), then the difference kinetics (Δ*W*_OK_ = *W*_OK_ (treatment) – *W*_OK_ (control)) were plotted in the linear time scale (Fig. 5A). Between O-step and K-step, hidden L-band (~100 μs) could be observed by such a subtraction. As researchers reported (Tsimilli-Michael, 2020), negative L-band was the signal of good energetic connectivity or grouping of PSII units, while positive L-band would be caused by stress. Therefore, figure 3A demonstrated that, L-band of ST plants could be observed clearly, and the value of Δ*W*_OK_ at L-band increased with time. On the contrary, RT plants remained negative L-band with slight fluctuation from 2 DAT to 8 DAT. In the same way, to explore whether the application of glyphosate lead to the emergence of K-band, raw fluorescence data were double normalized by *F*_0_ (20 μs) and *F*_J_ (2 ms), as *W*_OJ_ = (*F*_t_ – *F*_0_)/(*F*_J_ – *F*_0_), the difference between treatment and control (Δ*W*_OJ_ = *W*_OJ(treatment)_ – *W*_OJ(control)_) was calculated and plotted on a logarithmic time scale (Fig. 5B). The trend of curves was similar to Fig. 5A, K-band (~300 μs) in ST plants was elicited at 6 DAT and up to maximum at 8 DAT. However, RT plants were still below the zero level at all sampling points. It indicated that glyphosate affected the inactivation of the oxygen-evolving complex (OEC) centers in sensitive cultivars. Of course, this effect would get more serious with time. Thus, GS plants were more sensitive to glyphosate than GR plants due to poor resistant mechanism manifested in decreased energetic connectivity of PSII units and the inactivation of OEC.

**Fig. 5.**
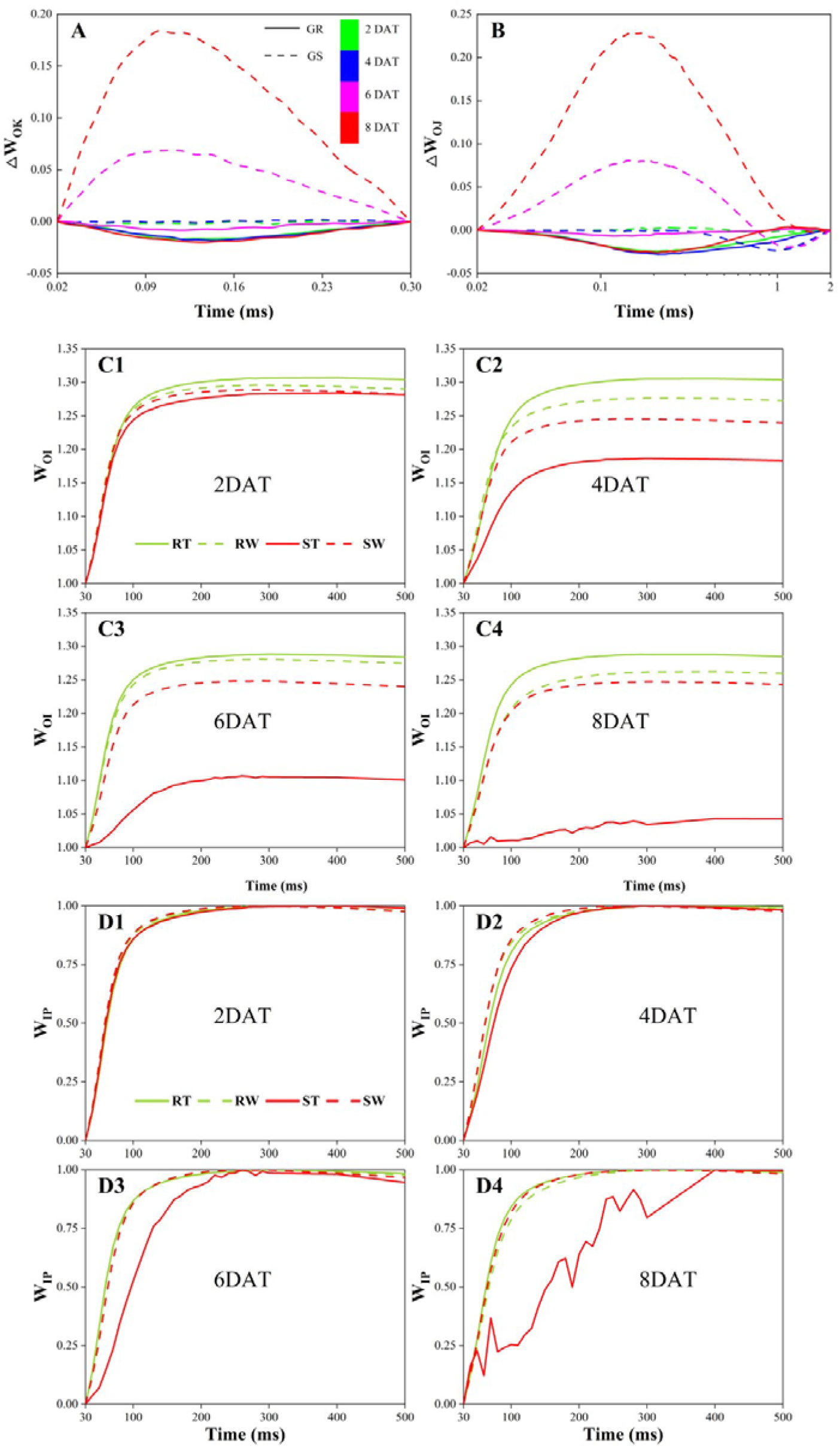
Changes in the (A) OK, (B) OI, (C) OP, (D) IP-phase of fluorescence rise kinetics of different glyphosatetolerance cultivars. *W*_XY_ = (*F*_t_ – *F*_0_)/(*F*_Y_ – *F*_X_), Δ*W*_XY_ = *W*_XY(treatment)_ – *W*_XY (control)_)

Besides, owing to the considerable information of PSI acceptor side, IP-phase was evaluated by two frequently used normalization procedures. As shown in Fig. 5C, double normalization between O-step (20 μs) and I-step (30 ms), as *W*_OI_ = (*F*_t_ – *F*_0_)/(*F*_I_ – *F*_0_), was plotted in the linear time scale (only *W*_OI_ > 1 was shown). In Fig. 5D, double normalization between I-step (30 ms) and the peak of fluorescence intensity (~500 ms), as *W*_IP_ = (*F*_t_ – *F*_0_)/(*F*_P_ – *F*_I_), was plotted in its linear time scale. Therefore, we could compare the changes between different cultivars/treatment/days after treatment from *W*_OI_ =f(t) and *W*_IP_ = f(t). The maximal amplitude of *W*_OI_ curve was positively related to the relative size of the end electron acceptors pool at PSI acceptor side. From Fig. 5C, RT plants had higher amplitude than RW plants while the opposite phenomenon was appeared in GS plants, which might be related to the resistance mechanism of glyphosate-resistant cultivar. Comparison of *W*_IP_ = f(t) could provide insight into the overall reduction rate of the end electron acceptors pool at PSI acceptor side, which rule out possible effects on pool size. Here it shown that glyphosate resulted in a decreased rate only in ST plants (Fig. 5D). Generally, the half-time, corresponding to *W*_IP_ = 0.5, was negatively correlated with electron transport rate. Thus, from the analysis of IP-phase, it could be concluded that glyphosate-introduced led to greater size of the end electron acceptors pool without change in electron transport rate for GR plants, while smaller size of the end electron acceptors pool and lower electron transport rate for GS plants.

### Effect of glyphosate stress on PSII and PSI by JIP-test

As a quantitative method, on the basis of several assumptions and approximations (Tsimilli-Michael, 2020), JIP-test plays an important role in the translation of raw fluorescence signals into biophysical parameters. Some functional and structural parameters were selected to explore the effect of glyphosate stress on PSII and PSI of different tolerance cultivars.

PCA was first applied to overview the entire ChlF data, and acquire a potential pattern of difference caused by glyphosate tolerance between wild and transgenic maize to stress in a more intuitive way (Fig. 6). The first two principal components explained 70.0%, 86.3%, 86.7%, and 92.7% of JIP-test parameters variation among cultivars under different treatment on 2, 4, 6, and 8 DAT, respectively. On 2 DAT, all groups could not be separated from each other. Since 4 DAT, four groups were divided into two categories well separated in score plot, where ST formed a distinct cluster, indicated there was a different reaction of ST from the other groups. Specifically, from 4 DAT, ST, positioned on the negative side of the PC1, were mostly determined by higher values of absorption and trapping per reaction center (ABS/RC and TR_0_/RC), relative variable fluorescence at J-step (*V*_J_) and at I-step (*V*_I_), and initial slope of normalized OJIP transient curve (*M*_0_). The other three groups (RT, RW, and SW), classified into the other category, were basically in the positive region of PC1 from 4 DAT onward. Apart from these, parameters δ_R0_ and ABS/CS_0_ devoted lower contribution to discrimination of different glyphosate-tolerance genotypes compared to other parameters.

**Fig. 6.**
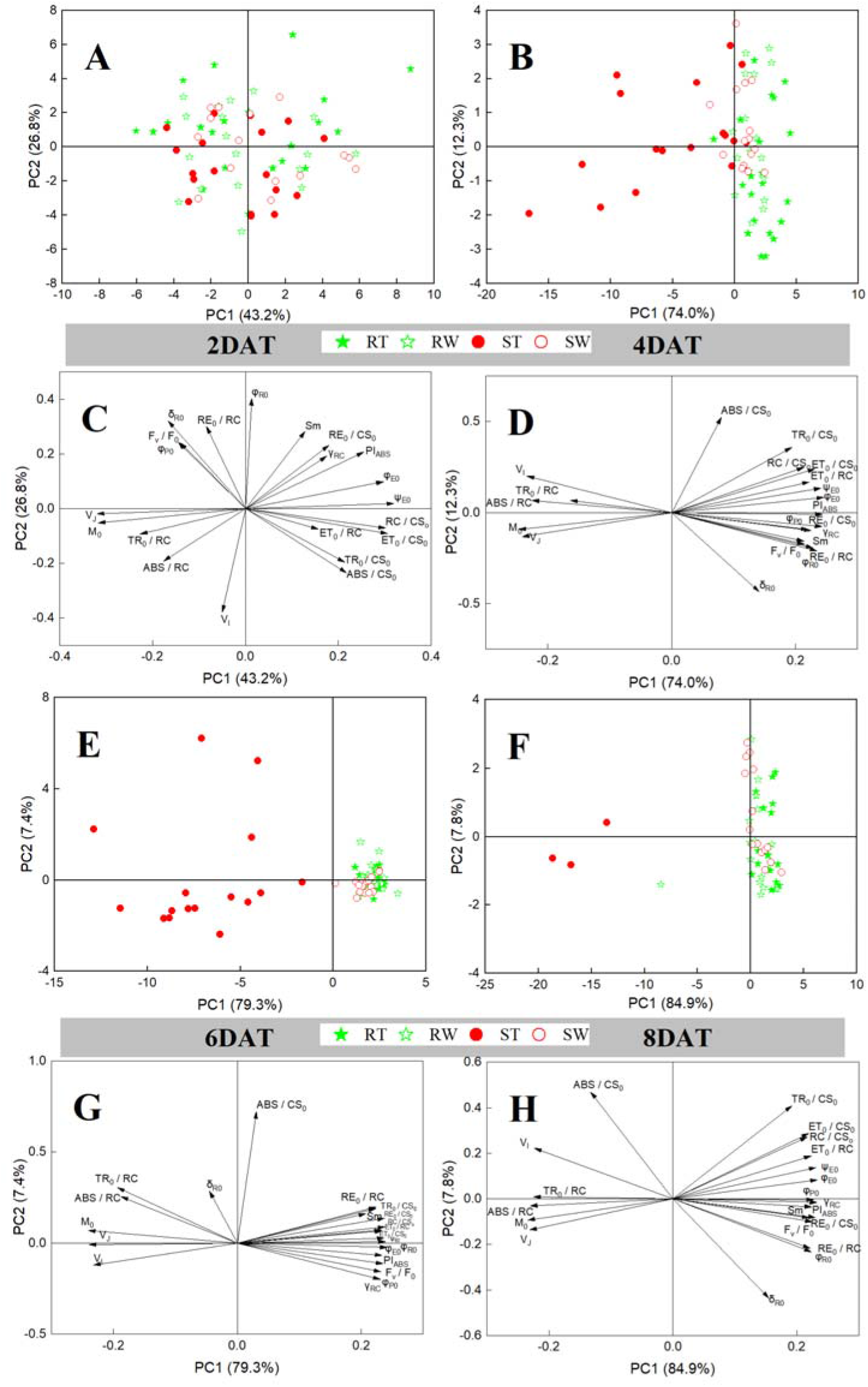
Principal component analysis of variation among 21 selected JIP-test parameters for RT (green solid pentacle), RW (green hollow pentacle), ST (red solid circle), SW (red hollow circle) on 2 (A, C), 4 (B, D), 6 (E, G), and 8 (F, H) days after treatment (DAT). Correlation between variables (C, D, G, H) and distribution of samples (A, B, E, F) along the first two PC axes are presented.

More specifically, all values of JIP-test parameters were presented as the ratio of glyphosate treatment to corresponding control on each sampling point for better readability._From the results of ANOVA (Table S2) and Fig. 7, most parameters of GS plants deviated from 1 (donated as the value of the control group). Parameters directly extracted from raw fluorescence data, such as ratio of original fluorescence data (*F*_v_/*F*_0_) and normalization complementary partial area above the OJIP curve on the maximal variable fluorescence (*S*_m_) were significantly decreased from 4 DAT and 6 DAT, respectively, while relative variable fluorescence at J-step (*V*_J_) and I-step (*V*_I_), and initial slope of the *V*_t_ = f(t) kinetics (*M*_0_) were notably increased along with time from 4 DAT. For the specific energy fluxes per Q_A_^-^reducing PSII reaction center, glyphosate decreased parameters ET_0_/RC and RE_0_/RC of GS plants significantly from 4 DAT, and improved ABS/RC and TR_0_/RC from 6DAT. Similar results could be found in the specific energy fluxes per excited cross section (CS) except for ABS/CS_0_, which did not change significantly until 8 DAT. As the index of RCs density, RC/CS_0_ and γ_RC_ were notably reduced from 6 DAT. Additionally, γ_RC_ also was used to estimate the ratio of chlorophyll in RCs and entire PSII. For quantum yields, four parameters (φ_P0_, ψ_E0_, φ_E0_, φ_R0_) were decreased significantly from 4 DAT in a whole. Besides, a strong decrease (about 44%) of PI_ABS_ could be found on 4 DAT, and another commonly used parameter, φ_P0_ (*F*_v_/*F*_M_) was also significantly decreased (about 8%) on 4 DAT. According to the degree of change, PI_ABS_ was more sensitive to glyphosate stress than φ_P0_ (*F*_v_/*F*_M_). Almost contrary to GS plants, all 21 parameters of GR plants were less affected by glyphosate, only *S*_m_ was notably decreased on 4 DAT.

**Fig. 7.**
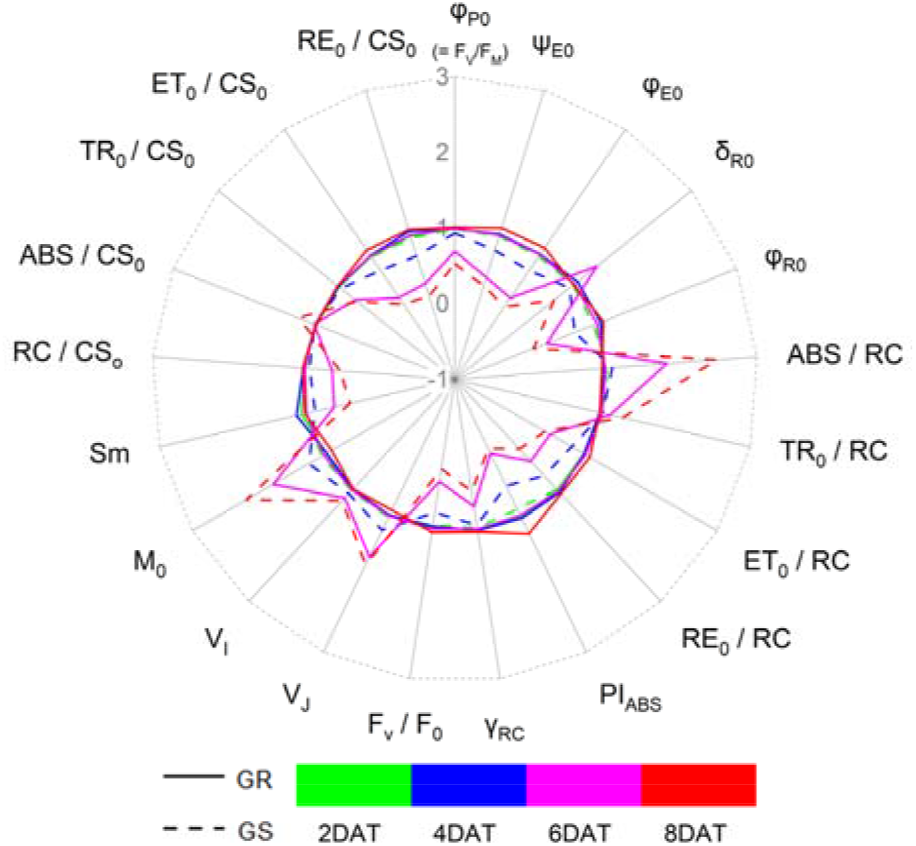
Relative changes of JIP-test parameters induced by glyphosate stress in maize seedlings (GR and GS) on 2, 4, 6, and 8 DAT. Taking each cultivar sprayed with water as the control (each parameter is set to 1, denoted by a black circle) and parameters of glyphosate treatment are expressed by fraction relative to the responding value of control. Individual sampling point is the mean value of replicates (3 times the number of plants).

### JIP-test parameters selection for responses of sensitive genotype to glyphosate

The above results suggested maize seedlings’ response to glyphosate is complex especially in time-scale. It is reasonable to select optimal JIP-test parameters in time-series manner instead of one particular day or all experimental period (Sun *et al.*, 2019). Therefore, PCA loadings was conducted to screen key features to evaluate glyphosate injury. According to the biplot (Fig. 6C, D, G, H), because the separation of ST and RT on PC1 was better than that of PC2 owing to higher explained variance, only the loadings of JIP-test parameters of PC1 was considered and sort by the absolute value of their loadings at each sampling point. The results were listed in order of importance (Table 2). From 2 DAT to 8 DAT, six selected features were not exactly the same, in which parameters φ_E0_, *V*_J_, ψ_E0_, and *M*_0_ had the highest frequency. To validate the performance of optimal features, cluster analysis was carried out on glyphosate-stressed plants (RT and ST) on each sampling point. Since the noteworthy shift occurred from 4 DAT to 6 DAT, only the Ward.D linkage clustering results of these days were shown (Fig. 8). Selected parameters exhibited the same pattern in PCA results (Fig. 6) using all parameters. With the advance of stress time, the separation degree of resistant and sensitive cultivars gradually increased, and on 6 DAT, RT and ST were distinctly categorized into two clusters indicating selected features could recognize plants of each genotype on 6 DAT. It could be concluded that the selected features explained most of the information of all JIP-test parameters, and contributed significantly to the response of photosynthetic performance pattern of maize seedlings under glyphosate-stress.

**Fig. 8.**
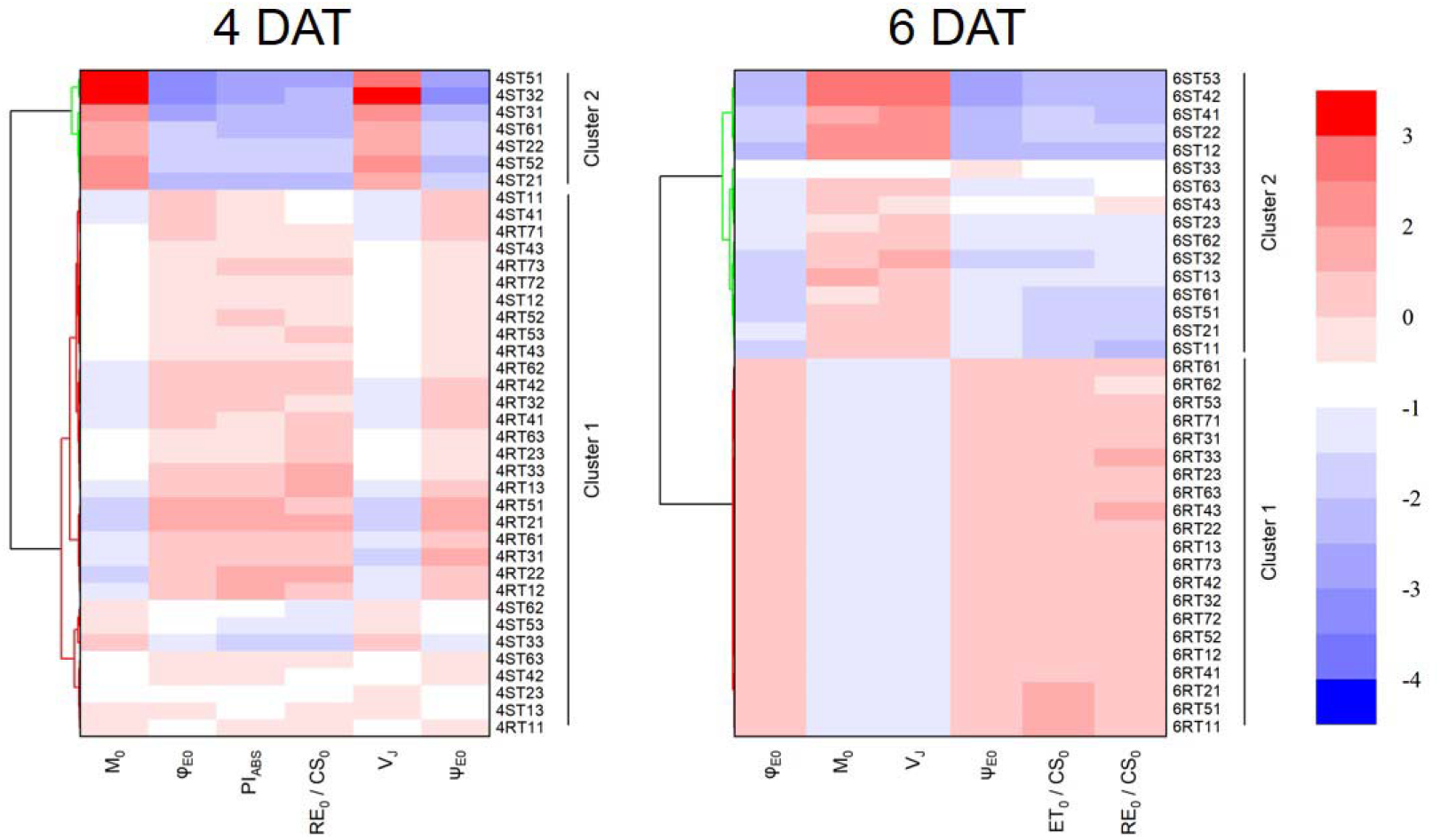
Clustering of samples using the Ward.D linkage algorithm with selected features of PCA loading on 4 and 6 days after treatment (DAT).

**Table 2.** Optimal features selected by PCA loading for each sampling day. They were listed in order of importance. DAT, days after treatment.

| **Sampling point** | **Selected JIP-test parameters** |
| --- | --- |
| 2 DAT | $V_J, \psi_{E0}, M_0, ET_0/CS_0, RC/CS_0, \varphi_{E0}$ |
| 4 DAT | $M_0, \varphi_{E0}, PI_{ABS}, RE_0/CS_0, V_J, \psi_{E0}$ |
| 6 DAT | $\varphi_{E0}, M_0, V_J, \psi_{E0}, ET_0/CS_0, RE_0/CS_0$ |
| 8 DAT | $M_0, \varphi_{E0}, \gamma_{RC}, V_J, \psi_{E0}, ABS/RC$ |

## Discussion

### Robustness of LCC prediction model

It is a common phenomenon that even though the parameters are the same each experiment, the model or methodology developed with the data from the first experiment is not applicable to the second experiment dataset, let alone the datasets obtained under a new filed condition. Consequently, when no sample in Exp.3 was introduced into the model development from the datasets of Exp.1 and Exp.2 and without TCA applied, the performance of predicting LCC was relatively poor. In view of this, this study proposed two strategies to narrow the difference between source domain and target domain to finally improve the model robustness with higher accuracy. The results were consistent with previous literature (Wan *et al.*, 2020). Moreover, based on the proposed model, we analyzed the changes of LCC under glyphosate stress according to the predicted values, and the conclusion were consistent with previous studies. Hence, the proposed method could be extended to predict LCC between different times’ datasets.

### Glyphosate treatment effects on photosynthetic physiology

The response of maize seedlings under glyphosate stress is consistent with previous researches (Feng *et al.*, 2018). Prompt ChlF transient curve is sensitive to both biotic and abiotic stress (Yusuf *et al.*, 2010; Magyar *et al.*, 2018). In this study, typical OJIP transient curve appeared in both RT and ST plants (Fig. 4). With glyphosate, the decrease in *F*_M_ in the two different glyphosate-tolerant maize plants could suggest the dissociation of LHC and the disruption of energetic connectivity, and this is an inherent protective mechanism to avoid light damage, over-reduction of the PQ, and damage of PSII (Oukarroum *et al.*, 2009). On 6 DAT, the resistance or activation of GR plants adaptive mechanism works, while the plant phenotyping difference of between ST and SW plants becomes larger and larger, which indicated GS plants has a more robust defense strategy against glyphosate.

When plants are exposed to stress for some days, rapid ChlF induction kinetics curve may exhibit other bands before J-step (Jiang *et al.*, 2008; Yusuf *et al.*, 2010). By double normalization of OJIP curve, L-band and K-band are observed in OJIP curve of ST plants. OJ-phase gives information of PSII, for example, L-band is an indicator of the energetic connectivity or grouping of antenna pigment and each PSII unit (Yusuf *et al.*, 2010; Mlinarić *et al.*, 2017), reflecting the stability of the system and the utilization of excitation energy between PSII units. In our research, the OJIP curve of ST plants showed a positive L-band, and the amplitude of it increased with time, which indicated glyphosate affects the connection between antenna pigment and PSII reaction center, or it may be a sign of interruption in energy transfer under light capture condition. According to the review (Tsimilli-Michael, 2020), the positive L-band means that ST plants has a worse connectivity in PSII units than SW plants, and less energy exchange, resulting in a partial loss of stability of PSII units. Once exposed to glyphosate stress, they become more vulnerable. However, GR with glyphosate-stress maintained negative value at L-band, indicating that glyphosate has little influence on the connection performance of antenna and PSII reaction center and excitation energy utilization of resistant cultivars. The presence of K-band is attributed to an imbalance in electron transfer from PSII donor side to the acceptor side and can be used to indicate the activity of OEC (Strasser, 1997; Yusuf *et al.*, 2010). As the rate of electron transfer from OEC to the secondary electron donor tyrosine residue (Y_Z_) slows, Y_Z_ becomes a stable oxidation state (Z^+^), more and more P680^+^ accumulates, resulting in less electron transfer to the primary electron acceptor Q_A_ (Strasser, 1997). In this study, the emergence of positive K-band indicates that the electron transfer from the PSII donor side to the receptor side of glyphosate-sensitive cultivar is inhibited under glyphosate stress due to dissociation of OEC. At this time, the electrons of secondary electron donor Y_Z_ can still flow to the PSII reaction center normally, resulting in the continuous accumulation of Z^+^, which leads to the increase of *W*_K_ at ~300 μs and K-band appears. To illustrate IP-phase response to glyphosate stress more comprehensively, the most common approach is to compare curves *W*_OI_ = f(t) and *W*_IP_ = f(t) simultaneously. In curve *W*_OI_ = f(t), the amplitude of ST was lower than that of SW, indicating the electron acceptor pool on the PSI acceptor side is relatively smaller. *W*_IP_ informs the relative reduction rate per reduction pool (Kalaji *et al.*, 2018a; Tsimilli-Michael, 2020). In this study, *W*_IP_ amplitude of ST plants decreased with time due to changes of the reduction rate of PSI acceptor side, while the *W*_IP_ amplitude of RT plants were basically the same as that of RW plants.

JIP-test is widely used to describe abiotic pressure on plants. On 4 DAT, some parameters of ST plants changed significantly compared with SW plants. As the commonest evaluation index of stress tolerance, PIAB_S_ exhibited better performance than φ_P0_ (or *F*_v_/*F*_m_), which is in agreement with finding of published study (Ambreen *et al.*, 2021). PIAB_S_ is obtained by multiplying three independent parameters (Table 1), it can evaluate the effect of stress on photosynthetic apparatus more comprehensively and accurately (Gururani *et al.*, 2018). Glyphosate increased *V*_J_ and *V*_I_ for GS plants, which indicated the poor ability of accept electrons of PQ pool lead the limited electron transfer ability of Q_A_ from acceptor side to Q_B_ while the electron transfer from pheo to Q_A_ generates Q_A_^-^ (Kalaji *et al.*, 2014). This eventually leads to a large accumulation of Q_A_^-^ and a decrease in the electron transfer efficiency in PSII (Öquist *et al.*, 1992). Similarly, *M*_0_ is another important indicator of electron transfer ability from Q_A_ to Q_B_, and the decrease of *M*_0_ agrees with the above inference.

The increase in ABS/RC in ST plants could suggest the proportion of active RCs decreases and the absorbed light energy increases, which is related to the inactivation of light RCs or the increase of antenna size. This finding could be additionally supported by a decrease in RC/CS_0_. In general, the inactivation of RCs is associated with photoinhibition in plants, which refers to the decline in photosynthetic function when the amount of light absorbed by plants exceeds the amount that can be utilized by the photosynthetic system. Once photoinhibition occurs, plants will actively close part of RCs and emit excess absorbed light energy in the form of heat (Derks *et al.*, 2015; Gururani *et al.*, 2015). In this study, the increase in ABS/RC in ST plants is much greater than the increase in TR_0_/RC and the decrease in φ_P0_. It can be sure that the mechanism to protect itself from glyphosate stress exists in sensitive cultivar, and some studies have shown that plants are more prone to light suppression or light damage under stress (Zeng *et al.*, 2021). However, the resistant cultivar does not show obvious light damage with glyphosate treatment, indicating that the resistant cultivar could flexibly adjust the antenna size and absorbed energy according to the external environment (Bano *et al.*, 2021), while the sensitive cultivar could not. Hence, there is a lot of excessive excitation energy in the PSII reaction center, thus destroying the activity of RCs and dissipating it as heat.

Generally speaking, the concentration of reactive oxygen species (ROS) in plants is too low to cause harm to plants, but once plants are in a certain stress environment, electron transfer and other processes are inhibited, oxygen molecules are reduced after gaining electrons, then a large number of ROS will be produced in various forms. In our study, glyphosate decreased the content of leaf chlorophyll in both GS and GR leaves. As suggested by previous literature (Gomes *et al.*, 2016), chlorophyll content degrades with the increase of ROS in plants leaves, leading to the decline of total chlorophyll content. The chlorophyll content in RT plants was slightly lower than that in RW plants on 2 DAT (the difference was not significant based on the result of ANOVA, *P*>0.05), but then gradually returned to normal level, while chlorophyll content in ST plants was significantly lower than that in SW from 6 DAT. This phenomenon indicates that resistant cultivar can avoid glyphosate’s negative effects to the greatest extent on the basis of its own genetic advantages, but such finite protection could not completely avoid glyphosate’s damage for sensitive cultivars, which can also be supported by the significant reduction of φ_P0_ (*F*_v_/*F*_M_, represents maximum quantum yield of PSII primary photochemistry in the dark-adapted state). Therefore, although GR plants are also effected by glyphosate, the negligible impact on the photosynthesis and oxidation process is greatly slighter than GS plants (Zobiole *et al.*, 2011).

### Potential applications and future prospect

Based on time series datasets, TCA_SVM regression model with source domain datasets updating strategy contributes to improving the robustness of the prediction of LCC in different time. Although it is proposed based on leaf reflectance in the laboratory, it also has considerable potential in the field from canopy scale to estimate chlorophyll content. In addition, the proposed method framework can be extended to other different crops and physiological biochemical indexes, not just maize or chlorophyll. Considering ChlF exhibits great performance in detecting glyphosate tolerance, more cultivars with different level of tolerance should be further investigated simultaneously to define tolerance criteria more accurately. Further work will focus on these and develop a decision-making platform for chlorophyll content prediction and glyphosate resistant cultivars identification. Furthermore, to accelerate the application of remote sensor equipment with various sensors in agriculture, it is imperative to develop a comprehensive crop spectral database.

## Conclusion

In this study, we demonstrated the feasibility of using the LCC predict results from regression models to analyze physiological responses to glyphosate stress. By updating data sets with new samples and applying TCA algorithm, the model performance can be improved effectively (R^2^_p_ = 0.84, and RMSE_p_ = 4.03). We also investigated the potential of using a time-series rapid ChlF transient to dynamically dissect the photosynthetic physiological response of different glyphosate-tolerance maize cultivars. Combining LCC with ChlF data, it could be concluded that the oxidative stress caused by glyphosate and the detrimental effects on photosynthesis are interconnected. For glyphosate-sensitive cultivars, inhibition of shikimic acid pathway would lead to changes in redox status and act on the metabolism of plants, mainly reflected in photosynthesis. Nevertheless, glyphosate-resistant cultivars could deal with excess ROS by activating the more powerful antioxidant system. Besides, according to the result of PCA loadings, φ_E0_, *V*_J_, ψ_E0_, and *M*_0_ could be used to indicate damage caused by glyphosate stress to differentiate resistant cultivars. This research was the first attempt to analyze the response of LCC to glyphosate by using the improved machine learning algorithms with transfer strategies based on datasets from different experiment times. Moreover, rapid ChlF transient technique exhibits potential for stress response monitoring in maize breeding programs and could facilitate superior genotypes screening and developing.

## Supplementary data

The following supplementary data are available at JXB online.

**Supplementary Table S1.** Leaf chlorophyll content (LCC) prediction results by support vector machine regression and random forest regression models development from multi-temporal spectral reflectance.

**Supplementary Table S2.** The response of JIP-test parameters to time-course effect. Values are presented as the means ± SD. Within a column, any two data with a common letter are not statistically significant according to ANOVA (Holm-Bonferroni post-hoc test, *P* > 0.05).

## Acknowledgements

This study was funded by the key projects of international scientific and technological innovation cooperation among governments under national key R&D plan (grant no.2019YFE0103800)

## Author contribution

XF and MT designed the research. MT, XB, YW, GY and JZ performed the experiments. MT, JZ and XF analyzed image data and constructed the models. MT and XF wrote the manuscript. YH provided suggestions on the experiment design and discussion sections. All authors read and approved the final version of the article.

## Data availability statement

The data supporting the findings of this study are available from the corresponding author (YH), upon request. Other data supporting the findings of this study are available within the paper and within its supplementary data published online.

## Abbreviations

ANOVA: analysis of variance
ChlF: chlorophyll a fluorescence
CS: cross section
DAT: days after treatment
GR: glyphosate-resistant
GS: glyphosate-sensitive
HSI: hyperspectral image
LCC: leaf chlorophyll content
LHC: light-harvesting complex
OEC: oxygen-evolving complex
PCA: principle component analysis
PQ: plastoquinone
RC: reaction center
RF: random forest
SVM: support vector machine
TCA: transfer component analysis

